# Barcoding populations of *Pseudomonas fluorescens* SBW25

**DOI:** 10.1101/2022.09.30.510243

**Authors:** Loukas Theodosiou, Andrew D. Farr, Paul B. Rainey

## Abstract

In recent years evolutionary biologists have developed increasing interest in the use of barcoding strategies to study eco-evolutionary dynamics of lineages within evolving populations and communities. Although barcoded populations can deliver unprecedented insight into evolutionary change, barcoding microbes presents specific technical challenges. Here, strategies are described for barcoding populations of the model bacterium *Pseudomonas fluorescens* SBW25, including the design and cloning of barcoded regions, preparation of libraries for amplicon sequencing, and quantification of resulting barcoded lineages. In so doing, we hope to aid the design and implementation of barcoding methodologies in a broad range of model and non-model organisms.

## Introduction

Barcoding individual microbial cells with chromosomally-integrated short, random sequences, is a powerful approach for obtaining high-resolution understanding of the eco-evolutionary dynamics that shape microbial populations (Blundell and Levy 2014; Levy et al. 2015; Cvijović et al. 2018; Ba et al. 2019). In brief, the technique involves introduction of a large number of random sequences into a single chromosomal region of a population of cells, such that all lineages are uniquely identifiable. A single PCR reaction allows amplification of the barcoded region, while DNA sequencing of the product reveals the frequency of each unique code. Changes in barcode frequencies allow inference of mutational events and fitness effects. For example, a cell that acquires a beneficial mutation will cause the frequency of the linked barcode to increase within the population. The rate of increase can be used to quantitate contributions to fitness.

When changes are measured over the course of hundreds of generations detailed understanding of eco-evolutionary dynamics can emerge (Levy et al. 2015; Jasinska et al. 2020; Acar et al. 2020; Boyer et al. 2021; Aggeli et al. 2021), including insight into the connection between genotype and phenotype of even complex traits (Venkataram et al. 2016; Kinsler et al. 2020; Jagdish and Nguyen Ba 2022; Nguyen Ba et al. 2022; Aggeli et al. 2022; Ascensao et al. 2022). Recent developments allowing regeneration of barcode diversity extend opportunity to observe evolutionary change over the course of thousands of generations (Ba et al. 2019).

While application of barcoding initially focussed on adaptive evolution in single populations of model microbes, particularly yeast (*Saccharomyces cerevisiae*) (Blundell and Levy 2014; Cvijović et al. 2018) and *Escherichia coli* (Jasinska et al. 2020), strategies have since been devised for studying the evolution of cancer using barcoded lung cancer cells in mice (Rogers et al. 2018) and quantification of eco-evolutionary dynamics in microbial communities (Venkataram et al. 2022). The potential for extending strategies and applications is considerable, ranging from tracing object provenance (Nguyen Ba et al. 2022), enhancing experimental power through mutant-multiplexing (Jackson et al. 2020), extending understanding of microbiome transmission (Vasquez et al. 2021), to the study of demographic changes during colonisation of new environments (Brettner et al. 2022)._ However one impediment to progress is the effort required to develop the initial barcoded library, especially in non-model systems.

Here, we outline a step-by-step guide for barcoding microbial cells, with particular focus on the bacterium *Pseudomonas fluorescens* SBW25 (hereafter SBW25). In addition to detailed protocols covering library preparation, amplicon sequencing methods and strategies for assessment of data quality, critical steps are emphasised and guidance of a general nature given in order to facilitate development of barcoded libraries in non-model microbes.

## Library creation

### A. Design and construction of barcoded integrative plasmids

The primary aim is to introduce barcoded sequences into the chromosome of an otherwise isogenic population in order to enable lineage-tracking in later experiments. Note that barcoding for lineage-tracking is distinct from ‘Tn-seq’ (van Opijnen et al. 2009), which introduces transposon insertion mutations along with a barcode sequence. The standard strategy behind barcoding for lineage-tracking involves creation of a vector with a region of extreme sequence variation, which is eventually introduced into the target bacterium. The backbone vector should have minimal fitness consequences and should integrate just once at a single genomic site. Additionally, the ‘barcode’ region should be short enough to be sequenced with short-read next-generation sequencing platforms. The downstream library of barcoded mutants should also be highly diverse (i.e., there should be many barcodes, each at a similar frequency).

For our libraries, derivatives of mini-Tn*7* vectors (Choi and Schweizer 2006) were employed. This transposon-based system ensures chromosomal integration of random (barcode) sequences in a single, known, chromosomal location (Waddell and Craig 1989). The strategy is similar to that used to barcode *E. coli* (Jasinska et al. 2020). The advantage of the Tn*7* system is the highly conserved position of the *att*Tn*7* site (downstream of the *glmS* gene) into which Tn*7* elements integrate (without causing inactivation of an open reading frame). Additionally, the *att*Tn*7* region is found across many species of interest (Mitra et al. 2010) and integration of Tn*7*-vector systems have been demonstrated across many organisms (Schlechter et al. 2018; Wiles et al. 2018), including SBW25 (Liu et al. 2014; Lind et al. 2015; Farr et al. 2017). In bacteria lacking an *att*Tn*7* site, such a site can be engineered into the genome, provided the bacterium of interest is capable of transformation and undergoing allelic replacement (Figueroa-Cuilan et al. 2016). If introduction of *att*Tn*7* is not an option, other methods are available for introducing barcodes, but they are less ideal for final applications. These options include: A) transformation of cells with barcoded plasmids (Vasquez et al. 2021), and B) the use of Tn*5*-based systems. The former requires selection for maintenance of the plasmid during subsequent evolution experiments and this can be problematic from the perspective of cost. The latter involves transposons that integrate effectively at random and thus stand to disrupt genes on insertion. This is clearly is highly undesirable.

In developing the integration vector for SBW25 all efforts were made to keep the vector as small as possible, and the metabolic cost of the vector as minimal as possible. The final vector is ~3000 bp and contains a minimal set of expressed genes. Limiting the size of the vector has the additional benefit of increasing the rate of transformation (although it should be noted that a co-transformation with a larger helper vector is required for integration of mini-Tn*7*-vectors). Our vector was assembled by Gibson assembly, although any assembly method can be used, including standard restriction enzyme digestion and ligation strategies.

The vector has three key elements: 1) an origin of replication (pBR322 origin) that allows propagation in an *E. coli* host (the pBR322 origin does not allow replication in non-*E. coli* hosts); 2) a selectable marker (in our case, a gene encoding tetracycline resistance: *tetA;* 3) paired Tn*7* integrative elements (L and R elements) between which the selectable marker and later the barcodes are situated (see Fig. 1A). Transcriptional terminators were included on either side of the selectable marker to prevent any unwanted expression of the antibiotic resistance gene, the downstream barcode, or the neighbour genes at the integration site.

**Figure 1:**
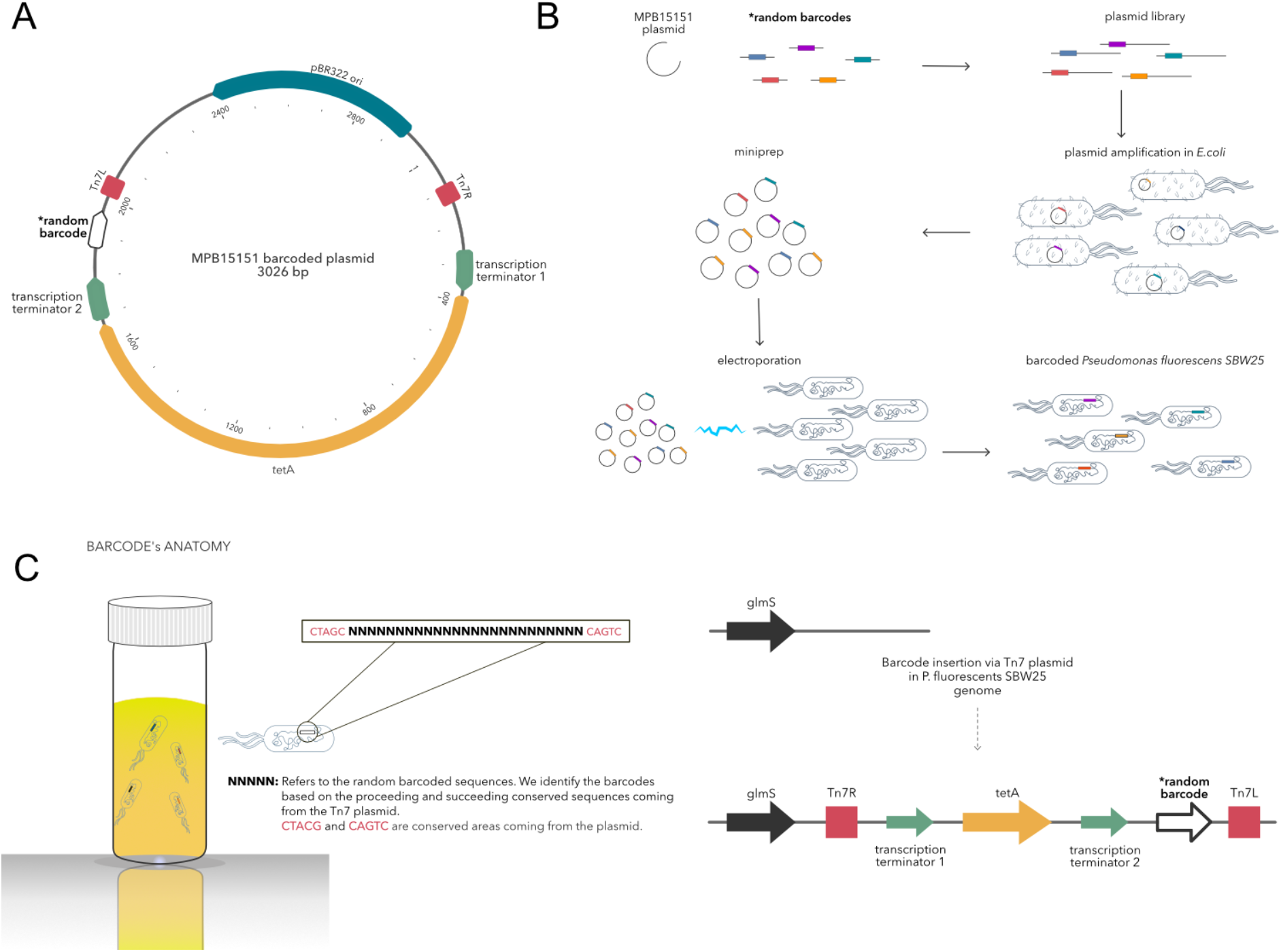
Barcoding *Pseudomonas fluorescens* SBW25 cells. **Panel A** shows a genetic map of the ~3 kb vector used to introduce barcoded sequences into the chromosome of SBW25. The vector features an origin of replication (pBR322) which allows replication solely in *E. coli* hosts, but not *Pseudomonas*, two *Tn7* integration elements that enable integration of intermediate DNA at the *att*Tn*7* site, a gene conferring tetracycline resistance(*tetA*) (allows selection for transformants), two transcription regulators to limit unwanted transcription of *tetA* and the barcode, and finally the barcoded region, consisting of 25 random base pairs. **Panel B** summarises the molecular process required to barcode SBW25. This process was repeated 25 times to maximise the diversity of the final barcoded library. Following the construction of the backbone vector, plasmids with 25N random sequences were used to PCR amplify the vector, the product was ligated into a circular vector and amplified by transformation into *E. coli* Top10 cells (transformants were selected on agar plates supplemented with tetracycline), a pool of plasmids were extracted from the resultant colonies and transformed into SBW25 with a pUX-BF13 helper vector (which expresses transposition machinery but does not replicate in SBW25). Transformants were grown on selective agar plates, and the resulting colonies were harvested and pooled into a single library. **Panel C** illustrates the 25 bp of a random barcode with 5 bp conserved areas left and right of the barcode, which can be used to extract barcode sequences from Illumina data. In addition, panel C shows the position downstream of *glmS* where the barcoded vector integrates into the SBW25 genome and final orientation of the vector.

Integration of random sequences (the barcodes) requires several rounds of cloning before the final barcoded library is delivered into the desired target species. During this process – detailed in supplementary files, visually summarised in Fig. 1B, and recounted in the following paragraphs – care should be taken to ensure the frequency of each barcoded genotype is as even as possible. Unevenness implies one or several barcodes are at a high frequency, which increases the likelihood that spontaneous mutations will occur in the lineage of those genotypes, reducing the resolution at which beneficial mutants can be quantified. To this end, repeated restriction digestion was used to limit ancestral vector sequences and the entire barcoding process was repeated 25 times so that overrepresented barcodes from any transformation would be reduced by 1/25.

To integrate random sequences into the previously described vector, the backbone was amplified with primers containing randomly generated 25 bp sequences (the “barcodes”) using a fast-cloning approach (Li et al. 2011). The PCR product was digested with DpnI to limit ancestral sequences, and the resulting products were transformed into electrocompetent TOP10 cells to both ligate the linear PCR product into a circular plasmid and amplify ligated plasmids. Transformed colonies from one plate were counted to estimate the total number of transformants. Remaining colonies were harvested, resuspended in LB and plasmids extracted over several miniprep columns. To improve the level of barcode diversity, the protocol was repeated 25 times. An estimated 580,000 transformant colonies were harvested for plasmid extraction at this step. The plasmids underwent a enzymatic digestion step to limit occurrence of the ancestral non-barcoded plasmid to <0.001% in the final barcoded population (note that without the final digestion, the ancestral barcode-free sequence was measured at about 1% of the final population of transformed SBW25).

Clonal populations of SBW25 were prepared for transformation with the barcoded vector using pre-established methods (Choi and Schweizer 2006). The barcoded plasmid and the pUX-BF13 helper plasmid (Bao et al. 1991) were transformed into SBW25 by electroporation. Cells from each transformation were plated on selective plates with a dilution series used to estimate the total number of transformants. Importantly, the total number of harvested SBW25 transformants was always less (~25%) than the number of *E. coli* transformants from which plasmids were extracted. Limiting the number of SBW25 transformants reduces the potential for transformation with the same barcoded vector, thus increasing diversity of the final library. The estimated number of unique barcodes was ~100,000. To further increase library diversity, the library-making process was repeated 25 times drawing upon plasmids from independent *E. coli* transformations. Because plates with more colonies have smaller colonies, measures of OD were made of resuspended cells with adjustments to dilutions to ensure equal contributions of each transformation to the final library of cells. Each of the 25 transformations was mixed in a final volume of ~50mL, which was then vortexed thoroughly and frozen for later experiments.

Following library construction, simple quality control checks should be made. The most important check involves determination of the fraction of clones that do not contain barcodes. Such unwanted cells – those containing no Tn*7* and those with Tn*7* but no barcode – can appear in the final library and arise as, for example, a consequence of imperfect antibiotic counter-selection or incomplete restriction digestion of the original plasmid. This fraction, if large, would mean that numerous mutational events are unassociated with a unique sequence and thus go undetected. This check was made by sequencing across the Tn*7* insertion site in 96 independent clones. The resultant data showed that all 96 contained unique barcodes.

### B. Library Preparation

Identification and quantification of barcoded lineages in populations requires amplicon sequencing. Our approach involved a two-step strategy for amplification: 1, amplification via PCR of the barcoded region (using primers specific to the barcoded region) and 2, amplification with Illumina-specific primers (this product was then sequenced on Illumina platforms, see Supplementary material: Amplicon-library preparation protocol). Table S6 lists primers used for the rounds of PCR. The first pair of primers feature sequencing primer binding sites, 8N randomers to detect amplification biases (these are optional), a smaller heterospacer (see below), and the barcode vector-specific sequence for annealing of the primers. The heterospacer consists of either none, one, or two extra nucleotides, with all three primers mixed to reduce saturation of the flow cell with signals from the same sequenced nucleotide. The second primer pair consists of the P5 or P7 flow cell binding regions, eight nucleotide indices to allow identification of the reads, and finally, a region complementary to the first set of primers to allow amplification of the initial PCR product. Together, this results in a ~310 bp product, with the barcoded region able to be sequenced from either end using 150 bp paired-end reactions.

At the point of PCR, it is necessary to be aware of problems arising from amplification of random barcode sequences – a problem that becomes more acute with each amplification cycle. It was observed during the design of the library-generation protocol that more than 20 PCR cycles of the barcoded region resulted in multiple large aberrant amplicons. This led to the design of a limited cycle approach, which involved extracting large quantities of DNA from the experimental samples (~1μg). This means that a large number of cells in the population are sampled and ultimately provides greater resolution on the frequency of each barcode. After testing several commercial kits, a chloroform-based DNA extraction method was employed because this maximised extraction of DNA from cell pellets (see Supplementary material: Salt DNA extraction protocol).

The DNA extracted from colonies was used as a template for the first round of PCR of 14 cycles using a high-fidelity DNA polymerase. This first reaction was performed in quadruplicate per biological sample. The four PCR products were then purified into a single concentrated sample, which was used as a template for the second round of PCR (of only 5 cycles). The final amplicon was then purified of remaining primers and primer-dimers with a magnet-based size selective kit. The purified product was then amplicon-sequenced using an Illumina MiSeq DNA Flex Library Prep Kit.

### C. Read processing and library assessment

Following construction of a barcode library, the size and diversity of the barcoded library needs assessment. This section describes a pipeline for the extraction of barcodes from sequencing data and then suggests an experimental protocol for testing reproducibility and quality of the barcoded library.

When acquiring sequence data, it is advisable to check the quality of the amplicon sequences by filtering reads with low quality to avoid erroneous results in downstream analysis. If paired-end data is acquired, it is suggested to merge both reads, keeping the reads with the highest quality. The user should check that most barcode sequences have the expected length and work only with barcodes close to the expected values. Moreover, it is best practice to map amplicon sequences to a reference sequence, ensuring that the amplicon sequences are in the same direction (sequences mcanight be reversed or be in the reverse complement state after merging). Finally, barcode sequences should be retrieved by extracting sequence data that maps within the coordinates of the reference barcode sequence. Alternatively, barcodes can be extracted based on the conserved areas left and right of the barcode (Figure 1, Panel C) as in the *bartender* pipeline (Zhao et al. 2018). Using this approach, all sequences must have the same direction as the reference.

To assess the quality and reproducibility of the barcoded SBW25 library, three independent overnight cultures were initiated from the frozen stock (inoculated with ~1% V/V). From each genomic DNA was extracted, resulting in three amplicon libraries for sequencing.

After paired-end sequencing with MiSeq, two fastq files for each sample were acquired, with every file containing approximately 1.5 million sequences. Each sequence was ~150bp long and included the barcoded area (~25bp) and a conserved area left and right of the barcode (~120bp). To assess read quality, fastQC version 0.11.8 and Trimmomatic version 0.39 (Bolger et al. 2014) were used to retain only reads with a quality above 30. Subsequently, paired-end reads were merged with PEAR version 0.9.8 (Zhang et al. 2014). All sequences were then mapped to a reference barcode sequence with Minimap2 version 2.24 (using default options) (Li 2018), which is essential for ensuring that all sequences are in the same direction and orientation. A custom-made pipeline was used (see Data Availability section), and the sequence reads that mapped to the reference barcode were extracted.

Read mapping revealed 179.374 unique barcoded sequences from replicate 1, 178.993 from replicate 2 and 159.870 from replicate 3. More than 98% of all barcodes in all replicates had the expected length of 25 ± 2bp (Figure 2) and could be used for downstream analysis. Pairwise comparison for all replicates was made to find common barcodes (Figure 3) and frequencies were compared using Pearson correlations (Figure 4). Barcode frequencies between replicates were highly similar, leading to an R=1 in all cases, indicating high reproducibility. Around 56% of the unique barcodes were found in all three replicates, although this can increase significantly if the barcodes are clustered based on their sequence similarity (the clustering of barcodes is not considered further in this article). It is thus possible to attribute differences in the number of unique barcodes in each replicate to two different factors: 1) PCR methods during library preparation and the sequencing process that may contain errors, leading to changes in the nucleotide sequence and, thus, to different unique barcode sequences; 2) Sampling effects: at each stage of culturing, sampling and amplicon sequencing, multiple bottlenecks exist that may alter final barcode diversity.

**Figure 2:**
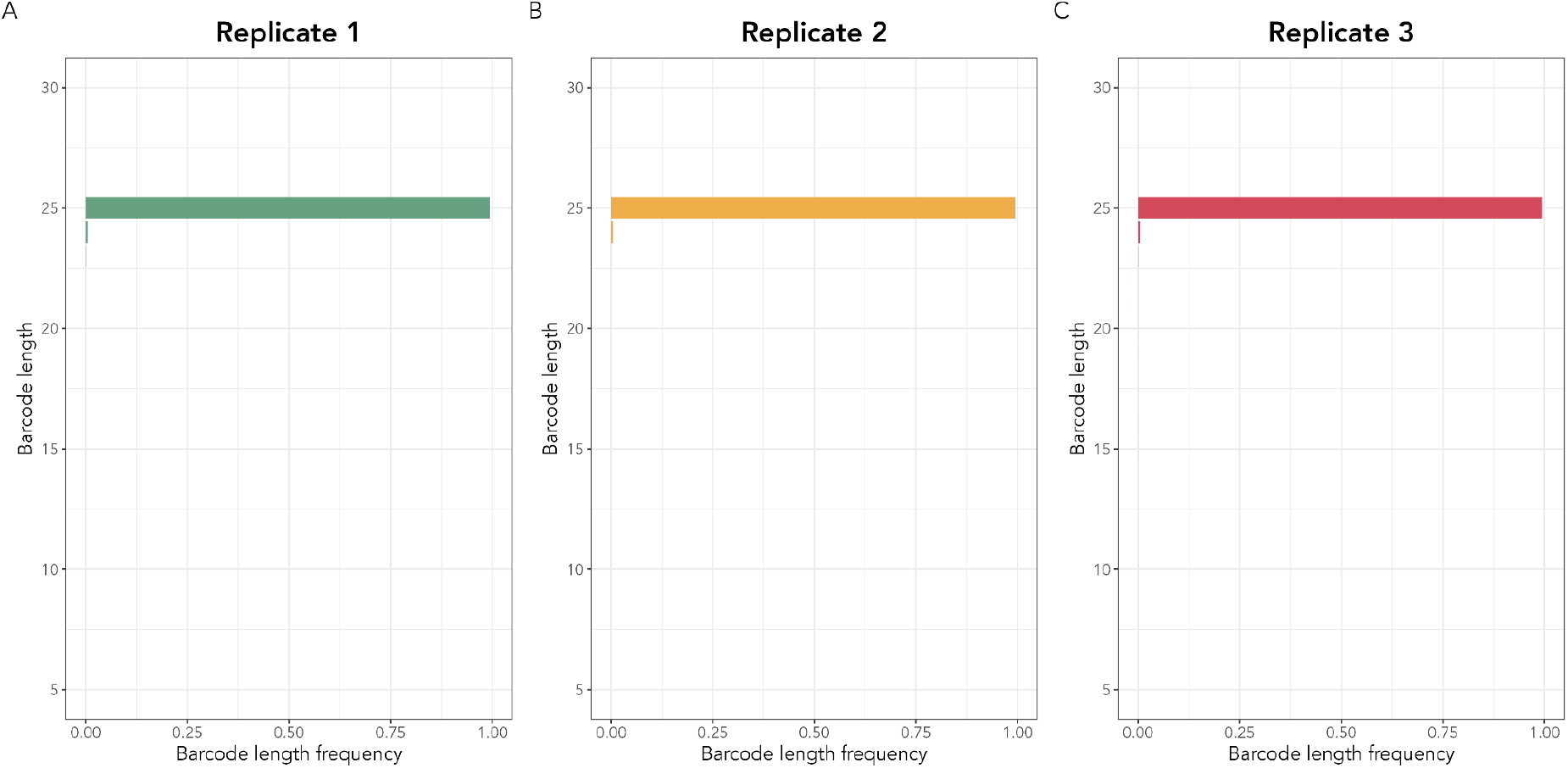
Frequency of barcodes with different lengths across replicates. The frequency of the length of all barcodes was estimated for each replicate. More than 98% of all barcodes in all replicates have an expected size of 25bp. Panel A indicates the frequency of barcode lengths in replicate 1, Panel B represents replicate 2 and Panel C replicate 3. The x-axis in each panel show the frequency of each barcode length, and y-axis indicates the barcode length.

**Figure 3:**
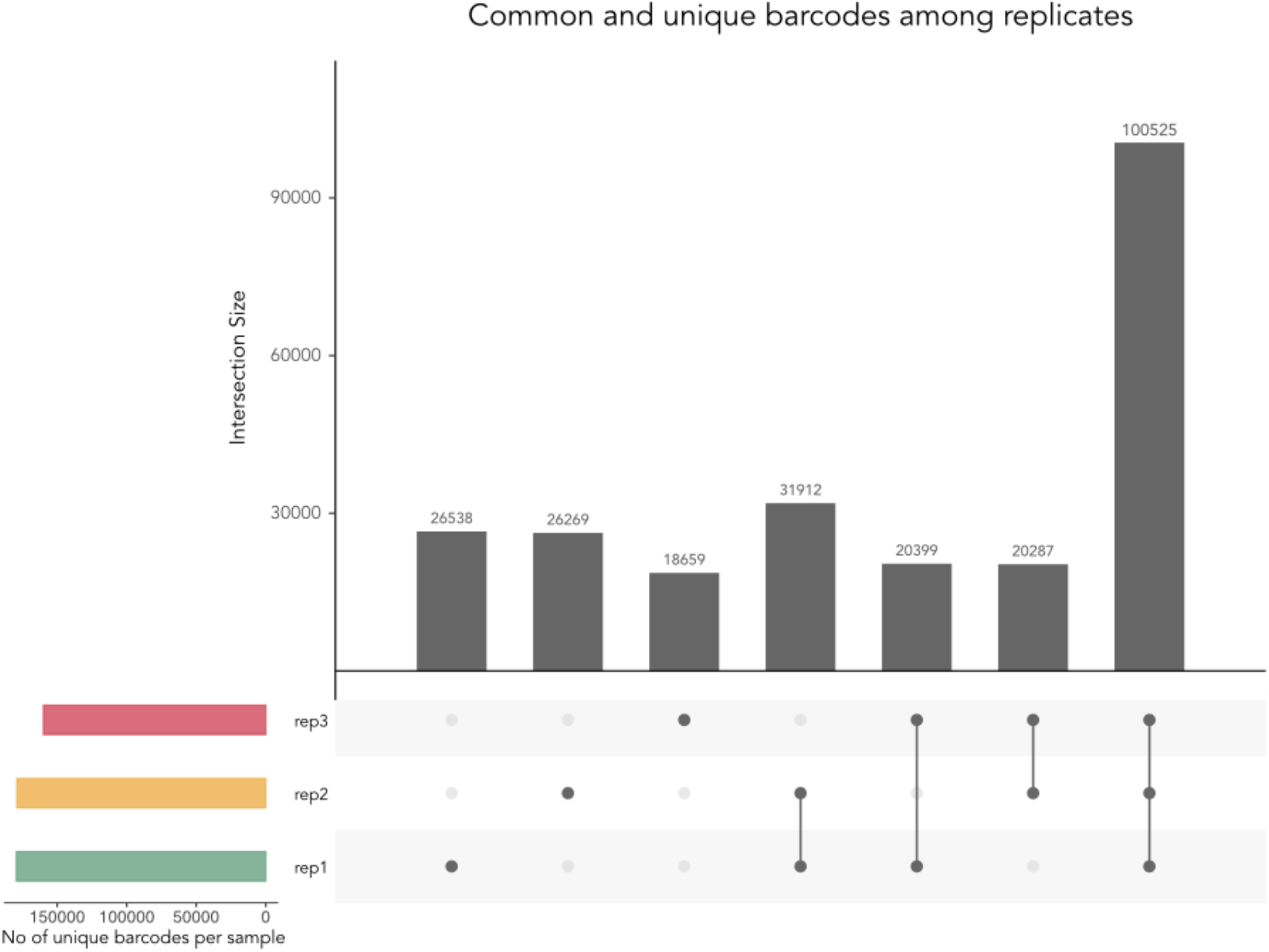
Common and unique barcodes among replicates. The UpSet plot shows the common and unique barcodes after comparing all sets of replicates. The coloured barplot at the lower-left part of the UpSet plot shows the number of unique raw barcode sequences. The dot matrix indicates the raw barcode reads in each set of replicates. Around ~55 % of all barcodes are common in all three replicates.

**Figure 4:**
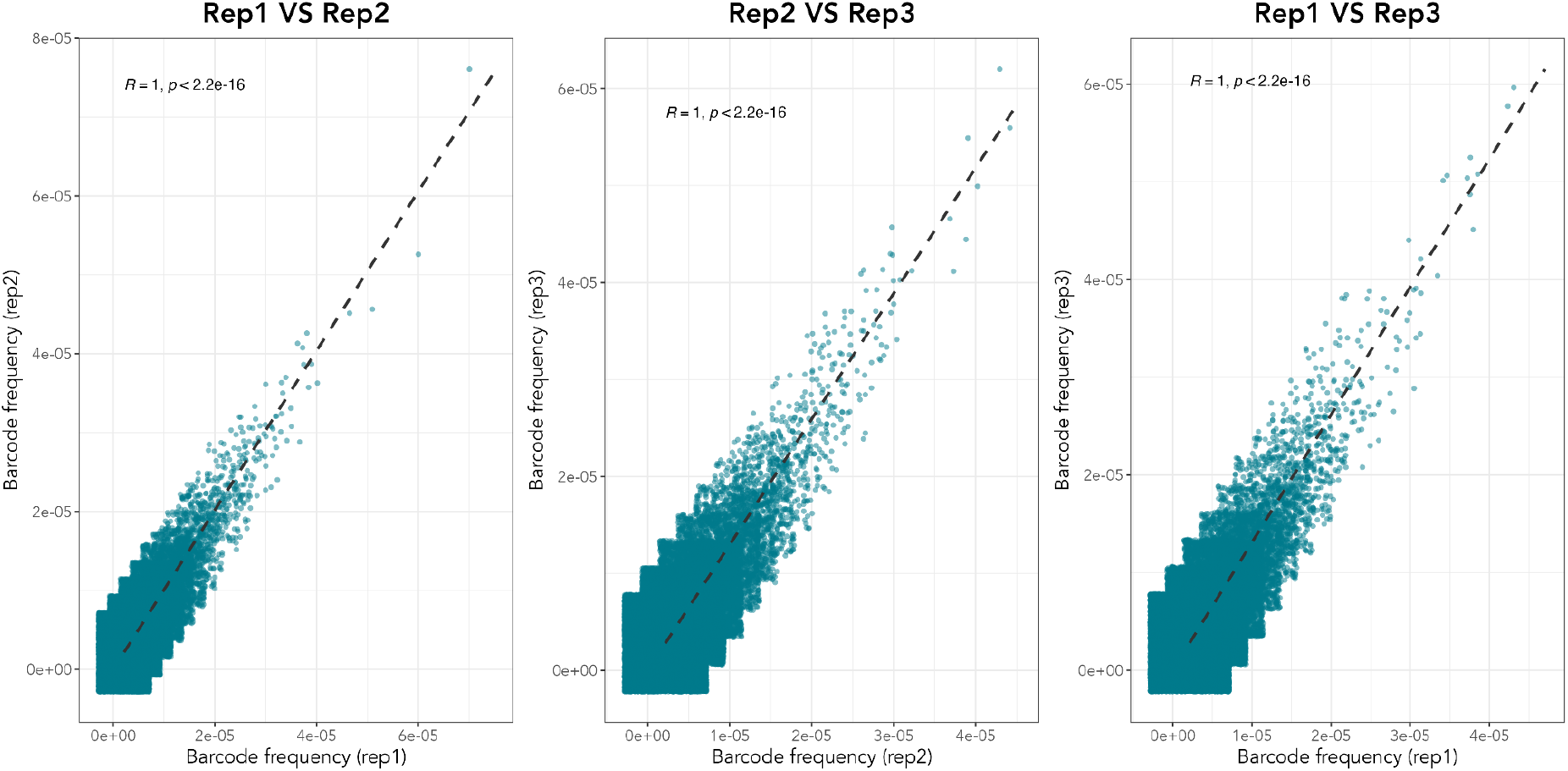
Pairwise comparisons of the barcode frequency between replicates. Following the extraction of the barcode reads, barcode frequencies between replicates were compared using Pearson correlation. Panel A compares replicates 1 and 2, Panel B compares replicates 2 and 3 and Panel C compares 3 and 4. Each panel indicates the R coefficient and *p-*value from the Pearson correlations at the top. The *R=1* in all panels coefficient indicates a high similarity of the barcode frequencies in all replicates.

### D. Next Stages

Having prepared and evaluated libraries, attention turns to experiments where barcoding can be applied. A standard experiment is to set up replicates of genetically homogeneous barcoded populations in different ecological conditions (e.g., with and without predators, antibiotics, salinity, or fluctuating environments), sample populations daily, amplify the barcoded region from the population and sequence the amplicons. The resulting sequences allow changes in the frequencies of lineages to be tracked. Aided by mathematical models, it is possible to infer the number of beneficial mutations, their fitness effects, and how these affect population dynamics of lineages. It is of interest to quantify competition among lineages and the contribution of ecology and evolution to lineage population dynamics.

Beyond analysis of adaptive evolution in single populations, barcoded organisms can be used to explore a range of questions in ecology and evolution. For example, introduction of a barcoded population to a community of microbial species allows analysis of the eco-evolutionary dynamics of the focal type embedded in a community (referred to as an ESENCE approach (Lenski 2017)). To do so, a population of the model or nonmodel organism can be barcoded and then introduced into experimental cultures composed of complex microbial communities. The dynamics of that focal strain can then be observed over time. Using this procedure, the effects of manipulation of the community (such as specific composition, diversity, resource use, or adaptation to resources etc.) can be assessed.

A second line of inquiry standing to benefit from barcoded populations concerns ecological factors shaping diversification in the face of dispersal and recolonisation events. Populations of barcoded cells can be subjected to different regimes of dispersal or disturbance over short time scales.

Once experimental data has been acquired, further challenges remain, such as clustering of sequencing data and inference of eco-evolutionary dynamics. The current methodology (Levy et al. 2015; Ba et al. 2019; Kinsler et al. 2020; Vasquez et al. 2021; Avecilla et al. 2022) for analysing barcoding data has been tailored to specific types of experiments that assume absence of ecological processes such as frequency-dependent selection. There is a considerable need to build pipelines for analysis that encompass more complex and realistic eco-evolutionary scenarios. The challenges presented require extensive development and description and will be the subject of a forthcoming paper.

### E. Conclusions

In this article we have provided a step-by-step guide for barcoding microbial cells: from library preparation to amplicon sequencing, including methods to assess the quality of sequence data. As an example, a protocol to barcode *Pseudomonas fluorescens* SBW25 has been provided, and procedures have been emphasised that enable maximum diversity of the resulting library. Barcoding SBW25, or any other strain, constitutes a powerful and versatile system for tracking the eco-evolutionary dynamics of populations – in a multitude of contexts – through time. We hope that the protocols described, along with accompanying discussion, serve to promote construction of barcoded libraries in a diverse array of organisms and development of new experimental possibilities.

## Acknowledgements

We thank Withe Derner for help constructing the barcoded cellular library, David Rogers for advice on the vector construction and amplification of the barcoded region, Ellen McConnell for her advice in improving the Salt DNA extraction protocol, Sven Kuenzel for his help sequencing the samples and Kaumudi Prabhakara for reading the manuscript and providing valuable insights.

## Data availability

All data and codes are deposited in the following publicly available GitLab repository https://gitlab.gwdg.de/loukas.theodosiou/sbw25-barcoding.

## Supplementary Material

### 1. Protocol for chromosomal integration of barcodes

Protocol to generate barcoded *Pseudomonas fluorescens* SBW25. This protocol starts from a specifically generated cloning vector (see MPB_15151_plasmid_unbarcoded in https://gitlab.gwdg.de/loukas.theodosiou/sbw25-barcoding) and details the introduction of the barcoded region and subsequent cloning of intermediate E coli hosts and the final SBW25 host. This process was repeated 25 times to maximise the diversity of the resulting library.

1. The plasmid of MPB15151 is extracted using the Miniprep Kit (Qiagen). The plasmid is digested with KpnI-HF (NEB) to linearise it following manufacturers’ instructions and purified using a PCR purification kit (Qiagen).
2. Using Primers MPB15151_BCSDM_25r4N_F and MPB15151_BCSDM_17HRF_R (see Table S1 for more details), 50ng of the digested plasmid was amplified using primers encoding the barcoded sequence in 20 μL volumes using ‘Q5 high-fidelity DNA polymerase’ (NEB) following the manufacturer’s instructions.
3. The entire PCR product was digested for 60 mins with the restriction enzyme DpnI (NEB) (following manufacturers’ instructions) and purified again with the PCR purification kit (Qiagen). DpnI digestion helps limit transformation with the ancestral plasmid.
4. 100ng (approx. 1 uL) of the digested amplicons were then transformed into ‘One Shot TOP10 Electrocomp *E. coli* (Invitrogen) using a ‘Micropulser’ electroporator (Bio-Rad), using a 0.1cm electrode gapped ‘Gene Pulser Cuvette’ (Bio-Rad). Cells were immediately revived with 500 μL SOC media in 1.5 mL microcentrifuge tubes, incubated at 37 °C and orbitally shaken at 1000 rpm.
5. The aliquot of revived cells was plated across 8 agar plates supplemented with 10μg/mL tetracycline, with one extra plating at a lower volume of cells (5 μL) to derive an estimate of transformants. Plates were incubated for 48 hrs.
6. Resulting colonies from each transformation were harvested by scraping directly off the plate and resuspended in 5 mL of LB supplemented with tetracycline (10 μg/mL)
7. 1mL from each resuspension was cryogenically stored, and the remaining 4mL was plasmid extracted using miniprep (Qiagen) - using 1 column per mL of cells - with columns eluted with 30 μL of EB warmed to 60 °C. Note at this stage, the plasmids are in their fully barcoded form (see the sequence of “MPB15151_barcoded_25N” in https://gitlab.gwdg.de/loukas.theodosiou/sbw25-barcoding).
8. Extracts from each miniprep were mixed, and 4 μg of plasmid were digested with KpnI-HF in 50uL volumes. This step limited the remaining ancestral vectors which do not contain the barcoded sequence (the barcode effectively replaces the KpnI restriction site). The KpnI-HF digested plasmid was purified using a PCR purification kit (NEB).
9. Electrocompetent *Pseudomonas fluorescens* SBW25 was prepared for electroporation using standard procedures (Choi and Schweizer 2006). To do so, overnight cultures were initiated from isogenic cryogenic stocks and grown over ~18 hrs at 28 °C. For each transformation, 1mL of overnight was washed over five cycles of centrifugation and washing with room temperature 300mM sucrose with the last resuspension made in 3μL of sucrose solution. Cells were kept at room temperature until electroporation.
10. Electrocompent cells were then co-transformed with 150ng of barcoded vector and the pUX-BF13 helper plasmid (Bao et al. 1991) using the same methods of electroporation and revival as the *E. coli* electroporation (see above). Cells from each transformation were plated on 5 LB agar plates (supplemented with 10 μg/mL tet) with an extra plate plated with diluted cells to estimate the number of transformants. Plates were incubated for 48 hrs at 28 °C.
11. Colonies were scraped together, resuspended in 3 mL of KB media, and vortexed for 40 secs. Colonies from the plate of diluted cells were counted to estimate the number of transformants harvested. Because plates with more colonies would have smaller colonies, measures of OD were taken of the resuspensions, and mild dilutions adjusted the volume of reach resuspension with KB to ensure equal contributions of each transformation to the final library of cells. As this step was performed on the same day, each of the 25 transformations was mixed in a final volume of ~50mL, which was then vortexed thoroughly, and frozen over approximately 50 aliquots with glycerol saline for later experiments.

### 2. Amplicon-library preparation

Protocol to generate indexed amplicons for 150 bp paired-end sequencing. Note: do not increase the number of cycles. A final amplicon product of 30 ng/μL is sufficient for Illumina sequencing.

1. Obtain high-concentration genomic DNA (approximately 500ng is used in this protocol for each PCR reaction).
2. Perform 1st PCR (Table S1)). Perform 4 PCR reactions (40 μL each) using DNA from the same sample. Use 500ng of DNA as a template in each 40 μL reaction. Add the Q5 polymerase last. The three primers composing Ampseq_HS0/1/2_F+24 and Ampseq_HS0/1/2_R-63 are mixed in equimolar ratios. Expect a final product of ~241 (The primers are ~64 and ~65 nucleotides long, and the template in between is 112 bp; for details, see table S1 and for the primer sequences, see Table S4).

**Table S1:**
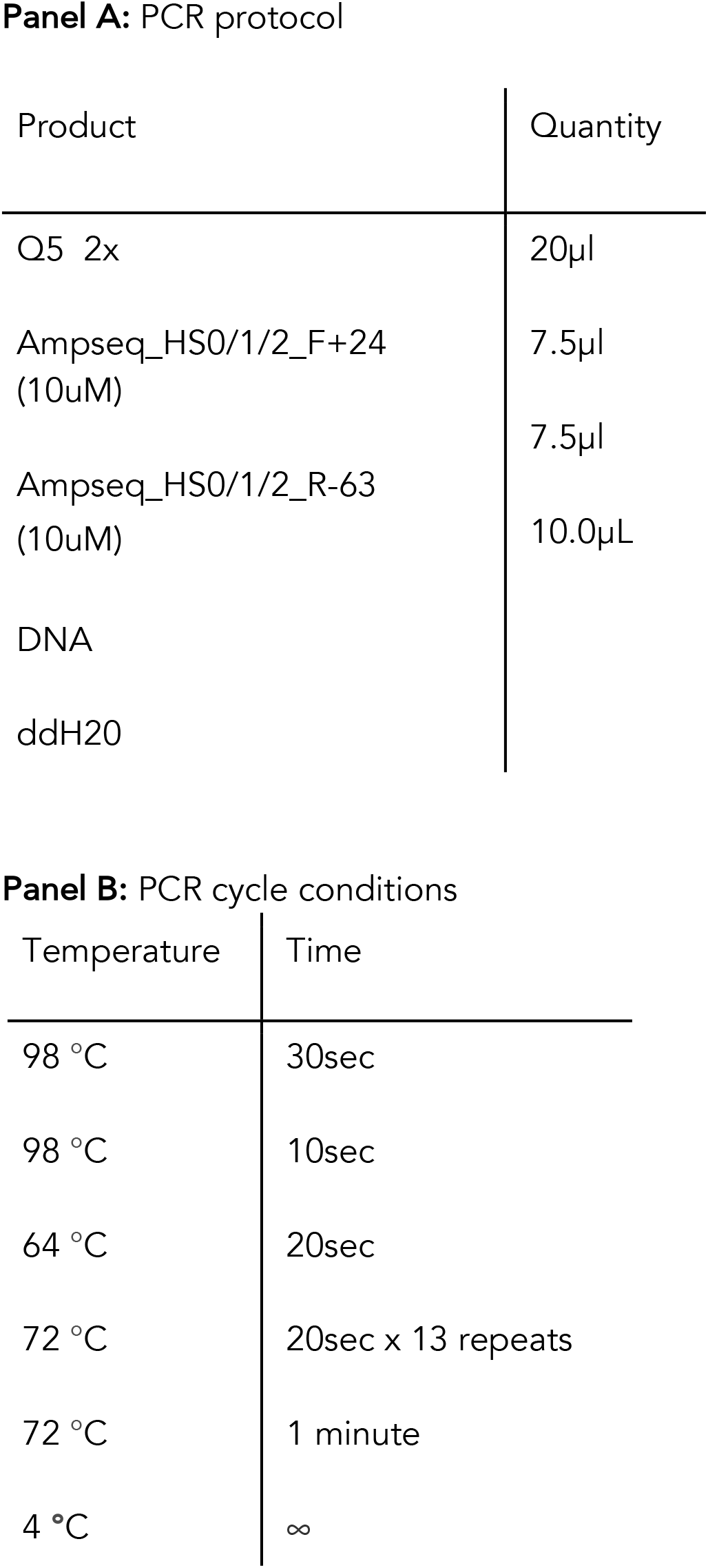
PCR protocol for the 1st PCR of the library preparation protocol. Panel A indicates the quantity of the PCR products that have been used, and Panel B indicates the PCR cycle conditions.
3. Gel electrophoresis is confirmed by amplicon (not a must at this point).
4. Remove genomic DNA and primers by DNA purification. Purify with Qiagen PCR purification kit - the main alteration being that the four PCR products are passed through one column when it is initially loaded. Elute each column with 40uL prewarmed H20. This is the template for the second PCR. A yield of about 5-50 ng/μL is expected at this stage.
5. Perform second PCR (Table S2) to add the second primer pair, which features the flow cell binding region and indices. You will use a lot more primer than usual to prevent template dimerisation. Expect a ~310bp product (the second primers are 37 and 32 nucleotides long, and the template in between is 241 nucleotides).

| Product | Quantity |
| --- | --- |
| Q5 2x | 20µl |
| Ampseq_HS0/1/2_F+24<br>(10uM) | 7.5µl |
| Ampseq_HS0/1/2_R-63<br>(10uM) | 7.5µl |
| DNA | 10.0µL |
| ddH2O |  |

**Panel B:** PCR cycle conditions
| Temperature | Time |
| --- | --- |
| 98 °C | 30sec |
| 98 °C | 10sec |
| 64 °C | 20sec |
| 72 °C | 20sec x 13 repeats |
| 72 °C | 1 minute |
| 4 °C | ∞ |

**Table S2:**
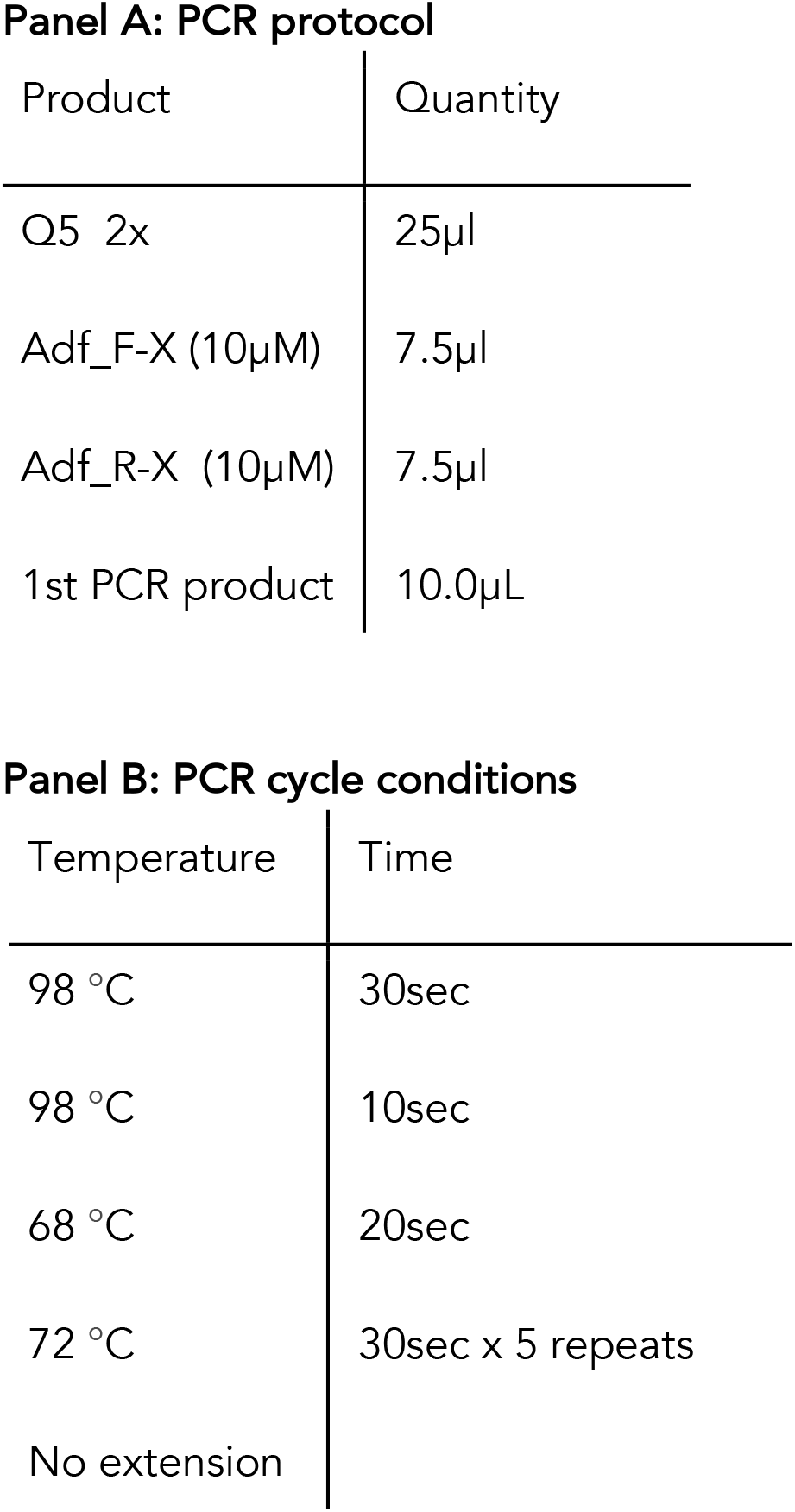
PCR protocol for the 2nd PCR of the library preparation protocol. Panel A indicates the quantity of the PCR products that have been used, and Panel B indicates the PCR cycle conditions.
6. Purify with Size selection to remove products <150bp (i.e. the primers must be removed) using a ‘ProNex Size-selective purification System’ (Promega), or similar size-selective strategy.
7. Run standard quality control methods before sequencing (check amplicon size ideally by automated electrophoresis to ensure the correct sized amplicon. Establish DNA concentration by fluorometry.

### 3. Protocol for DNA extraction of bacterial cultures

1. Spin the culture down (14500xg) and remove the supernatant.
2. Add 500μL HOM-Buffer (HOM-Buffer = 80mM EDTA, 100mM Tris, 0,5% SDS) and 5μL RNase. Note: We used the RNase A Solution 4mg/ml from Promega
3. Incubate samples for 1 h, at 37°C in a tube shaker.
4. Add 5μL Proteinase K [0,20mg/ml]. Note: We used QIAGEN Proteinase K ready-to-use solution 20 mg/ml
5. Incubate samples at 55°C overnight (500rpm) in a tube shaker.
6. Add 500μL of Sodium-Chloride solution (4,5M) and place samples for 10min at 4°C.
7. Add 300μl Chloroform and mix gently for 15min in a rotating mixer. Note: Chemicals such as chloroform should be used carefully under a hood.
8. Spin down for 10min at 10000rpm.
9. Transfer the upper phase (ca. 850μL) into a new tube.
10. Add 595μL of 100% isopropanol and mix for 5min in a rotating mixer.
11. Spin down for 10min at 13000rpm and remove supernatant.
12. Add 500μL 70% Ethanol (dilution from 100% ethanol for molecular biology) and incubate for 5min at room temperature. Note: We used Ethanol absolute Molecular biology grade from AppliChem
13. Spin down for 10min at 13000rpm and remove supernatant.
14. Repeat the last step.
15. Dry the pellet at room temperature.
16. Add 30μL of elution buffer and let samples stay for 30 min at room temperature.
17. Place the samples overnight at 4°C.
18. Measure DNA with Nanodrop and Qubit.
19. Store samples at −20°C.

### 4. List of primers that have been used in amplicon library preparation

All the primers that have been used for the barcoded library of SBW25 were HPLC purified, except of the initial fast-cloning primer encoding the random barcodes, which was PAGE purified.

**Table S3:**
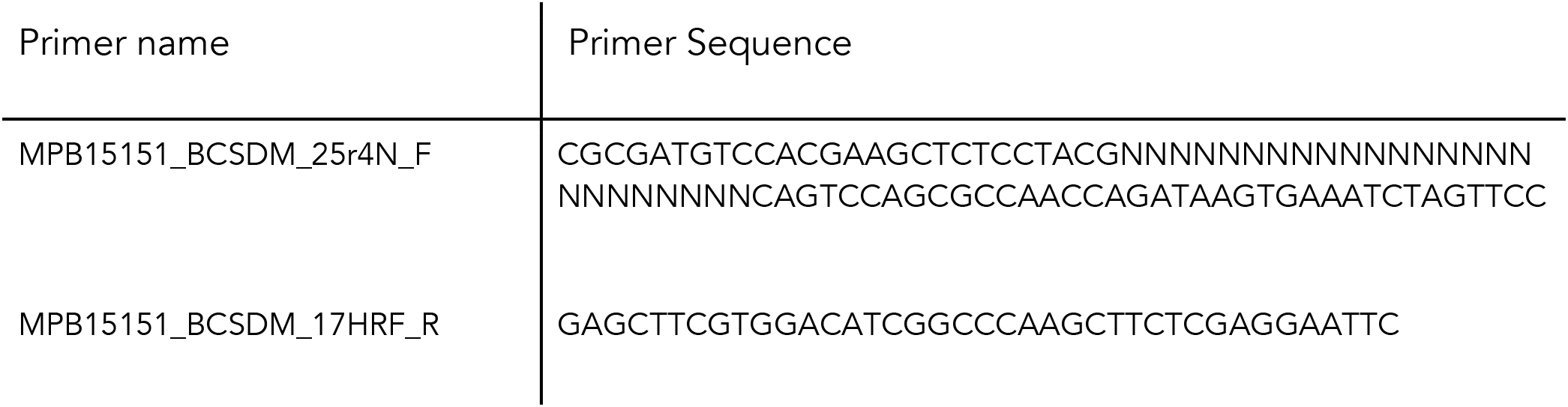
Primers that have been used for cloning.

**Table S4:**
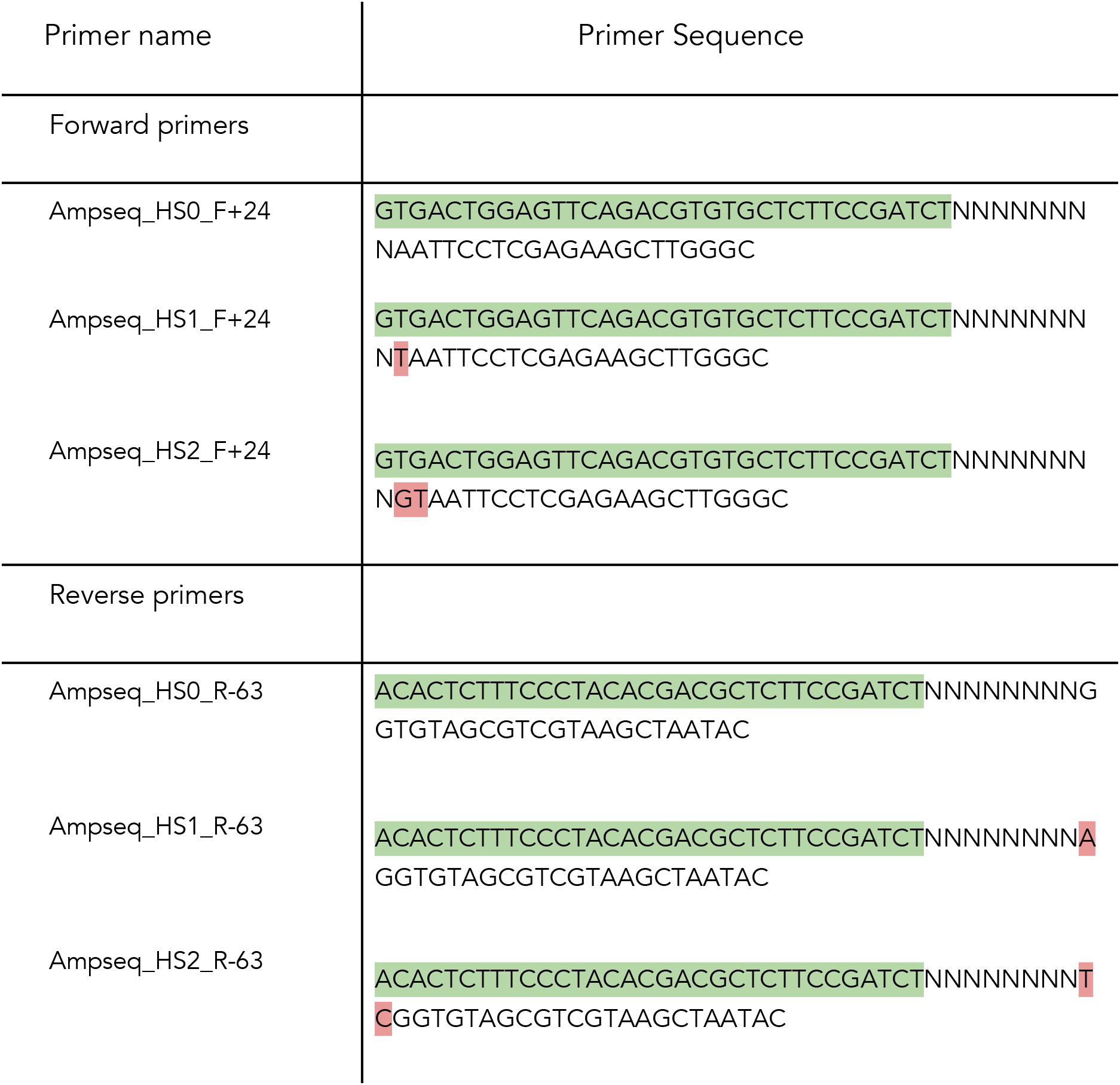
A list of the primer sequences were used for the 1 st PCR of the library preparation for sequencing. The green colour at the 5’ site indicates an Illumina sequencing primer binding region, the *8N* a random sequence, and the red colour is an heterospacer. Lastly, the non-coloured area at 3’ site indicates an *insitu* binding region.

**Table S5:**
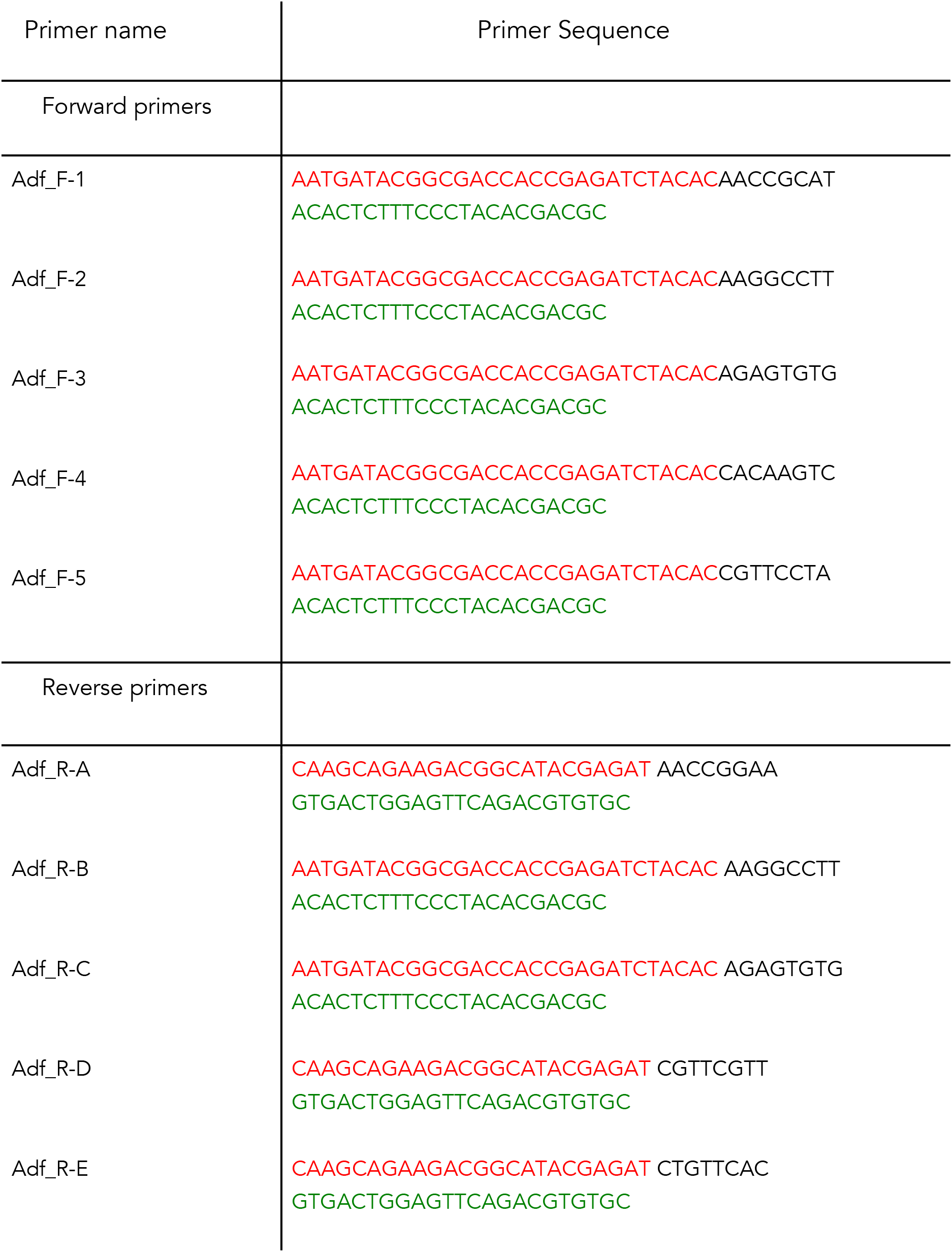
A list of the primer sequences used for the 2nd PCR of the library preparation for sequencing. The red text at the 5’ site indicates the P5 or P7 flow cell binding regions, while the black text, *8N*, depicts a random sequence. Finally, the green text at the 3’ site indicates a binding region for 1st PCR round.

**Table S6:**
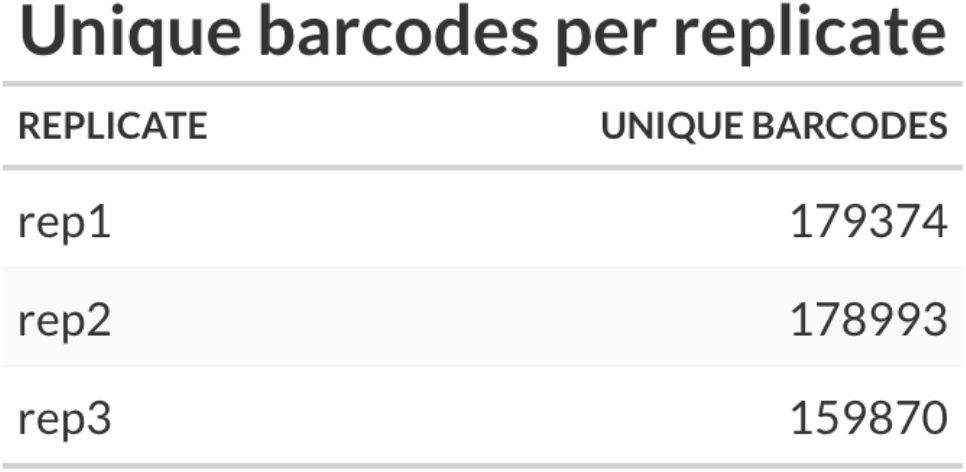
Number of unique raw barcode reads in each replicate.

Unique barcodes per replicate
| REPLICATE | UNIQUE BARCODES |
| --- | --- |
| rep1 | 179374 |
| rep2 | 178993 |
| rep3 | 159870 |

**Figure S1:**
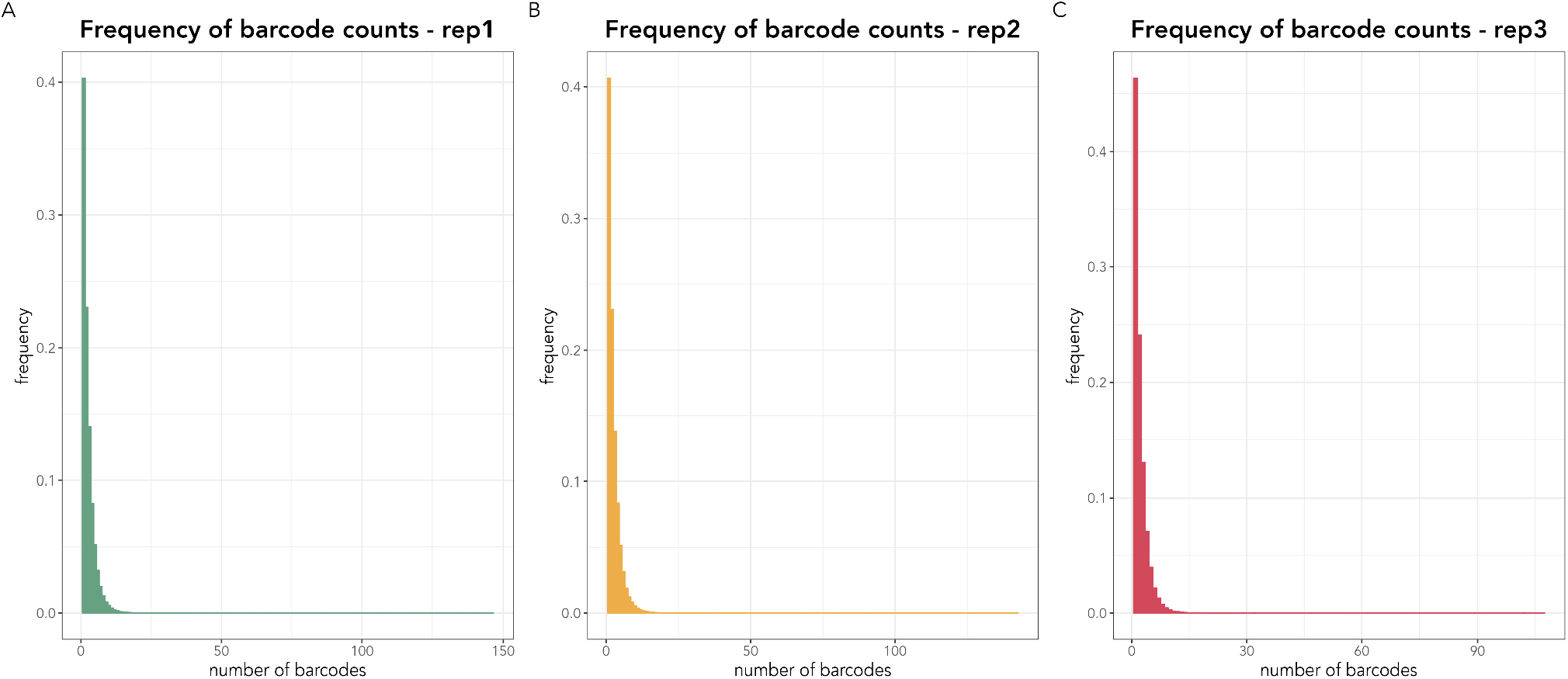
Frequency histograms of the barcode counts. A frequency histogram of the barcode counts for each replicate is presented in each panel. Panel A refers to replicate 1, Panel B to replicate 2 and Panel C to replicate 3. The frequency histograms are highly similar across replicates with the vast majority of the barcodes being at similar low frequencies.

## Notes

### Competing Interest Statement

The authors have declared no competing interest.

